# Data Descriptor: Human whole exome genotype data for Alzheimer’s Disease

**DOI:** 10.1101/2022.10.11.511653

**Authors:** Yuk Yee Leung, Adam C Naj, Yi-Fan Chou, Otto Valladares, Nicholas Wheeler, Honghuang Lin, Prabhakaran Gangadharan, Liming Qu, Kaylyn Clark, Laura Cantwell, Heather Issen, the Alzheimer’s Disease Sequencing Project, Sudha Seshadri, Zoran Brkanac, Carlos Cruchaga, Margaret Pericak-Vance, Richard P. Mayeux, Amanda B Kuzma, Wan-Ping Lee, William S. Bush, Anita Destefano, Eden Martin, Gerard D. Schellenberg, Li-San Wang

## Abstract

Bigger sample size can help to identify new genetic variants contributing to an increased risk of developing Alzheimer’s disease. However, the heterogeneity of the whole-exome sequencing (WES) data generation methods presents a challenge to a joint analysis. Here we present a bioinformatics strategy for joint calling 20,504 WES samples collected across nine studies and sequenced using ten different capture kits in fourteen sequencing centers in the Alzheimer’s Disease Sequencing Project. gVCFs of samples were joint-called by the Genome Center for Alzheimer’s Disease into a single VCF, containing only positions within the union of capture kits. The VCF was then processed using specific strategies to account for the batch effects arising from the use of different capture kits from different studies.

We identified 8.2 million autosomal variants. 96.82% of the variants are high-quality, and are located in 28,579 Ensembl transcripts. 41% of the variants are intronic and 15% are missense variants. 1.8% of the variants are with CADD>30.

Our new strategy for processing these diversely generated WES samples has shown to generate high-quality data. The improved ability to combine data sequenced in different batches benefits the whole genomics research community. The WES data are accessible to the scientific community via https://dss.niagads.org/.

## Background

The Alzheimer’s Disease Sequencing Project (ADSP) was established in 2012 as a key initiative to meet the goals of the National Alzheimer’s Project Act (NAPA): to prevent and effectively treat Alzheimer’s Disease (AD) by 2025. Developed jointly by the National Institute on Aging (NIA) and the National Human Genome Research Institute (NHGRI), the aims of the ADSP are to (1) identify protective genomic variants in older adults at risk for AD; (2) identify new risk variants among AD cases; and (3) examine these factors in multi-ethnic populations to identify therapeutic targets for disease prevention.

The ADSP completed whole-exome sequencing (WES) of 10,836 unrelated cases and controls in 2018. The data were generated by three NHGRI-funded Sequencing Centers (Broad Institute, the Baylor College of Medicine’s Human Genome Sequencing Center, and Washington University’s McDonnell Genome Institute) using Illumina technology and QC’ed by the ADSP. The study identified a novel rare variant in the long non-coding RNA *AC099552.4*, as well as two novel genes (*OPRL1* and *GAS2L2*), via gene-based analysis (Bis et al. 2020) that were associated with AD. However, the discovery of novel rare variants for AD is still limited by available sample size.

ADSP has sought to leverage other WES data sets (most of which were generated concurrently with the ADSP’s data set in the collaborative network) to increase the power to detect AD-related rare variants, not only limited to the Non-Hispanic White population but in other populations as well (African American, Hispanic, etc.). This collaboration will lead to the generation of the largest yet AD WES data set sharable with the public community. Combining data sets generated in projects that are originally designed for studying AD or other related dementias (ADRD) from different labs across different times (2010-2021) poses new challenges, as each WES data set was generated and processed using different protocols, potentially introducing biases into the combined data set (Clark et al. 2011; Sulonen et al. 2011; JS et al. 2011). As sequencing cost decreases and technology advances, more sequence data will be available shortly (both whole-genome sequencing [WGS] and WES) and will be generated using different protocols. Due to the desire for joint and meta-analysis, there is a need to process data generated by different platforms efficiently in a consistent manner.

To ensure all sequence data are processed following best practices with consistency and efficiency, the Genome Center for Alzheimer’s Disease (GCAD) in collaboration with the ADSP developed the genomic variant calling pipeline and data management tool for ADSP, VCPA (Leung et al. 2019). This is functionally equivalent to the CCDG and TOPMed pipelines (Regier et al. 2018). VCPA has currently been adopted as the official pipeline for processing all ADSP WGS/WES data, as well as data received from the collaborative network, a group of principal investigators (PIs) who have obtained either NIH funding or funding from private foundations involved in sequencing small numbers of AD samples.

Compared to WGS data, WES data focuses on exons, which make up roughly 1% of the entire genome. The critical challenge unique to WES data harmonization is using different capture kits to sequence samples. The capture kits were made by different vendors over the years containing probes that were designed using different reference genomes and versions of gene annotations.

In this study, GCAD built a new computational framework to integrate information from multiple capture kits (see Table 1 for details) while calling variants at the individual level and joint genotyping across individuals. Corresponding updated QC strategies were specifically developed for this data. Finally, all the individual data and joint-called data were shared with the community via https://dss.niagads.org/ in February 2020.

**Table 1:**
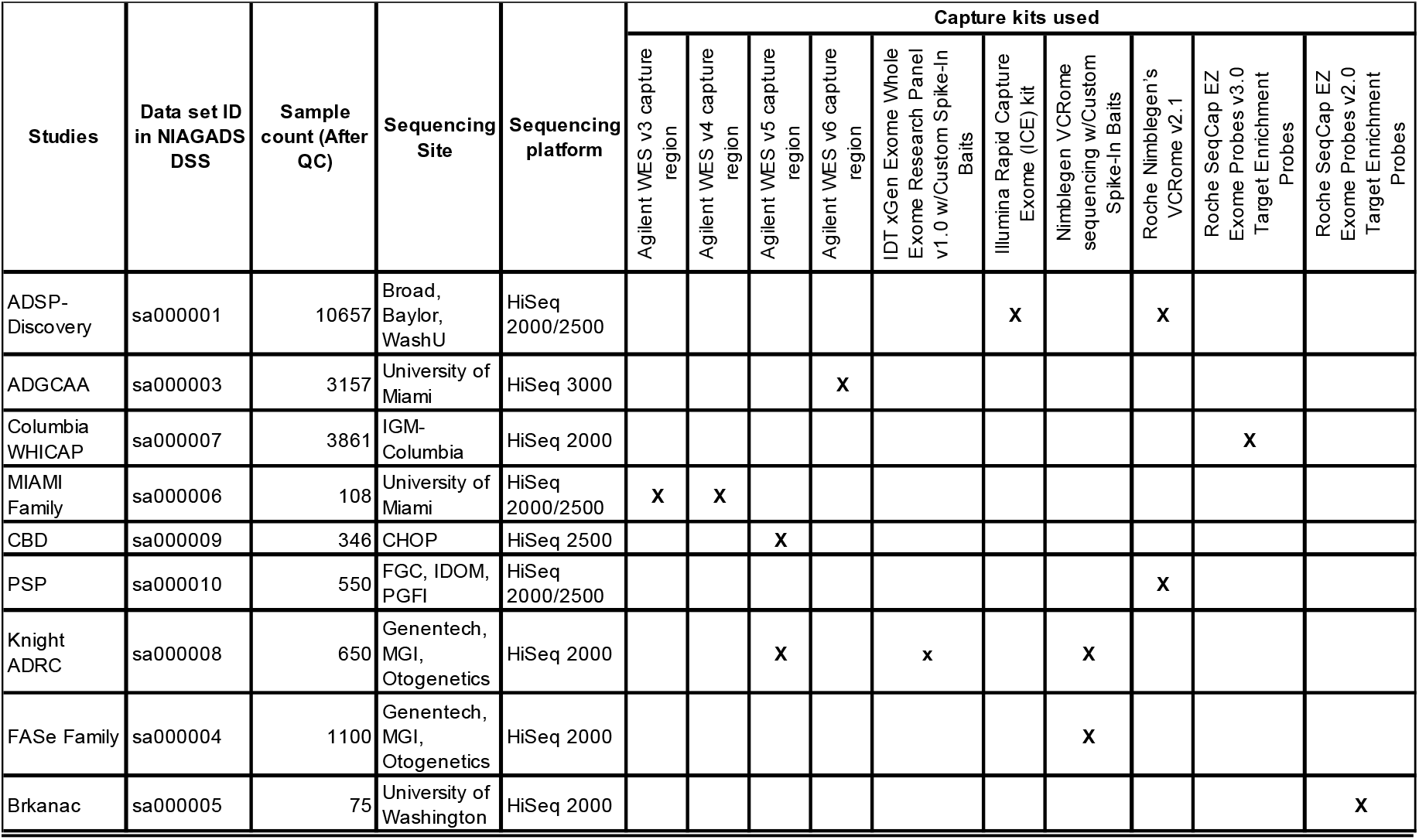
Summary of the data set with 20,504 WES samples from the nine studies, sequenced using ten different capture kits across fourteen sequencing centers. Footnotes for the sequencing site: the Broad Institute (Broad), Baylor College of Medicine Human Genome Sequencing Center (Baylor), the McDonnell Genome Institute at Washington University (WashU), Institute for Genomic Medicine - Columbia University (IGM-Columbia), Children’s Hospital of Philadelphia (CHOP), Functional Genomics Core of the Institute for Diabetes, Obesity and Metabolism (FGC, IDOM), Penn Genome Frontiers Institute - University of Pennsylvania (PGFI), Genentech company, (Genentech), MGI company (MGI), Otogenetics Corporation (Otogenetics).

## Data description

### Sample selection

The data set consists of 20,504 samples across nine studies (Table 1). Approximately half of the samples are from the ADSP-Discovery (part of the ADSP case/control data). Case control status were defined using the NINCDS-ADRDA (National Institute of Neurological and Communicative Disorders and Stroke, and the Alzheimer’s Disease and related Disorders Association) criteria (Beecham et al. 2017). GCAD reached out to ADGC/ADSP PIs in the collaborative network and received 9,847 additional WES samples from eight different studies. Table 1 contains Sample counts that have passed through QC (see **Section Sample level quality assurance checks** for details).

### Genome Sequencing and Capture kit information

Libraries were constructed from sample DNA with PCR amplification. Sequencing was performed across fourteen sites using different combinations of Illumina sequencing platforms and capture kits. Several studies (e.g., ADSP-Discovery) used multiple kits. while other studies used a single capture kit for all samples in their study design (e.g., Roche Nimblegen’s VCRome v2 kit was used in ADSP-Discovery, PSP, and Knight ADRC studies). The details are summarized in Table 1.

### Demographics

Phenotype information, such as disease status (AD and other dementia case or control), self-reported race/ethnicity, sex and age, as well as the number of APOE e2/e3/e4 alleles per individual, were obtained from the phenotype data shared by the data contributors. The demographics of this dataset are summarized in Table 2. This dataset contains subjects from three major populations: 13,362 Non-Hispanic White, 4,103 Black or African American, and 2,195 Hispanic. Altogether, there are a total of 9,955 cases (75.8 ± 8.9 years old) and 9,717 controls (81.5 ± 8.1 years old). 60% of the cases were female, with a similar gender ratio for controls. 44% of the cases (and 23% of the controls) have ≥1 APOE e4 alleles, which is a known genetic risk factor for AD.

**Table 2:** Summary of the demographics for each study. Listed in this table are total number of samples. Duplicate samples from the same subject for platform comparison are counted multiple times.

| Studies |  | Race/Ethnicity |  | Age |  | Gender (% of Female) |  | # of APOE e4 alleles |  |  |  |  |  |
| --- | --- | --- | --- | --- | --- | --- | --- | --- | --- | --- | --- | --- | --- |
|  |  | Cases | Controls | Cases | Controls | Cases | Controls | Cases |  |  | Controls |  |  |
|  |  |  |  |  |  |  |  | 0 | 1 | 2 | 0 | 1 | 2 |
| ADSP_Discovery | Non-Hispanic White | 5596 | 4279 | 75.6 (8.7) | 86.8 (3.7) | 58% | 59% | 57% | 40% | 3% | 87% | 13% | 0% |
|  | Hispanic | 234 | 157 | 75.2 (7.3) | 74.8 (8.4) | 65% | 59% | 59% | 40% | 1% | 61% | 39% | NA |
|  | Other/Unknown | 3 | 3 | 76.3 (11.8) | 86.3 (2.1) | 67% | 0% | 67% | 33% | NA | 100% | NA | NA |
| ADGC_AA | Non-Hispanic White | 1 | 0 | 58.0 (0) | NA | 0% | NA | NA | 100% | NA | NA | NA | NA |
|  | Hispanic | 2 | 6 | 81.0 (0) | 74.0 (12.0) | 100% | 100% | 50% | 50% | NA | 67% | 33% | NA |
|  | Black or African American | 1283 | 1634 | 74.5 (8.0) | 72.9 (8.2) | 70% | 74% | 35% | 49% | 15% | 62% | 35% | 3% |
| Columbia_WHICAP | Non-Hispanic White | 83 | 800 | 85.4 (5.1) | 80.5 (6.6) | 66% | 58% | 80% | 18% | 2% | 77% | 21% | 2% |
|  | Hispanic | 511 | 1257 | 84.0 (5.5) | 80.6 (6.3) | 75% | 70% | 69% | 29% | 2% | 79% | 20% | 1% |
|  | Black or African American | 218 | 939 | 84.0 (5.7) | 80.1 (6.6) | 76% | 69% | 61% | 33% | 5% | 67% | 31% | 2% |
| Miami_Families | Non-Hispanic White | 86 | 18 | 74.2 (7.6) | 76.6 (6.9) | 66% | 39% | 44% | 53% | 2% | 78% | 22% | NA |
| CBD | Non-Hispanic White | 335 | 0 | 63.5 (8.4) | NA | 45% | NA | NA | NA | NA | NA | NA | NA |
| PSP | Non-Hispanic White | 550 | 0 | 69.2 (8.4) | NA | 45% | NA | NA | NA | NA | NA | NA | NA |
| KnightADRC | Non-Hispanic White | 224 | 338 | 68.3 (8.8) | 71.3 (8.9) | 42% | 59% | 48% | 42% | 9% | 67% | 29% | 3% |
|  | Hispanic | 0 | 3 | NA | 78 (19.1) | NA | 100% | NA | NA | NA | 100% | NA | NA |
|  | Black or African American | 26 | 3 | 64.0 (9.2) | 74.3 (5.5) | 65% | 33% | 19% | 50% | 23% | 67% | 33% | NA |
|  | Other/Unknown | 3 | 2 | 76.0 (14.1) | 76.0 (9.9) | 100% | 0% | 33% | NA | 67% | 50% | 50% | NA |
| FASe_Families | Non-Hispanic White | 731 | 274 | 78.5 (7.2) | 78.8 (7.5) | 63% | 57% | 25% | 57% | 18% | 52% | 44% | 4% |
|  | Hispanic | 7 | 2 | 79.4 (3.3) | 75.0 (4.2) | 71% | 50% | 71% | 14% | 14% | 50% | 50% | NA |
|  | Other/Unknown | 2 | 2 | 72.5 (3.5) | 68.0 (4.2) | 100% | 0% | 50% | 50% | NA | NA | 100% | NA |
| Brkanac_Families | Non-Hispanic White | 44 | 0 | 74.9 (7.6) | NA | 61% | NA | 30% | 39% | 25% | NA | NA | NA |
|  | Hispanic | 16 | 0 | 68.1 (11.9) | NA | 56% | NA | 63% | 31% | 6% | NA | NA | NA |
| Total |  | 9955 | 9717 | 75.8 (8.9) | 81.5 (8.1) | 60% | 64% | 47% | 38% | 6% | 77% | 22% | 1% |

## Methods

### Integrating Multiple WES Capture kits

As summarized in Table 1, ten different capture kits were used for sequencing samples across nine studies. These capture kits were manufactured over the years by three different vendors (Illumina, Agilent, and Roche) and had substantial differences in kit contents, as the capture kits were generated based on different genomic annotation databases (e.g., Ensembl (Aken et al. 2016) on different reference genome builds [GRCh36, GRCh37]). Whenever possible, capture kit annotation files (in BED format) were received directly from the data contributors. If not, GCAD downloaded the original capture kit annotation files from the vendor’s website. Note that these files contain the genomic coordinate information, and do not include the exact sequences of the designed regions. If GRCh38 information of capture kits is not available, we performed UCSC liftOver (Kuhn, Haussler, and James Kent 2013) converting coordinates to GRCh38. All regions were further combined as a single BED file that contains the union of the capture kits’ genomic intervals (with flanking ± 7 bps). The BED files for individual capture kits, as well as the BED file containing all the captures’ intervals, are available in NIAGADS DSS (https://dss.niagads.org/datasets/ng00067/).

### Processing WES using VCPA at the individual sample level

VCPA, a BWA/GATK-based pipeline (DePristo et al. 2011; Li 2013), was developed by GCAD and the ADSP and optimized for processing large-scale, short-read WGS data (Leung et al. 2019). To adopt VCPA for ADSP WES data processing, GCAD followed GATK Best Practices (Van der Auwera et al. 2013) with the following additional steps to accommodate for the use of multiple WES capture kits. Instead of calling variants limited to the capture regions per sample (JS et al. 2011; Sulonen et al. 2011), VCPA keeps all detected variants, as we envision that (1) the research community will use the joint-called VCF for different kinds of analyses (i.e., one project will select a few studies for its analyses, but another project might pick different studies); and (2) Any WES data sets in the future may use different capture kits. Compared to VCPA-WGS, VCPA-WES has explicit steps as follows:

- Coverage calculation – the 20x coverage metric (i.e., percentage of base pairs [bps] with 20 reads or more) was calculated on regions that were included in the capture kits only.
- Variant evaluation – Ti/Tv ratio (ratio of the number of transitions to the number of transversions) and counts of SNPs/indels were calculated on regions that were included in the capture kits only.
- Variant calling – VCPA-WES has been updated to use GATK4.1.1 HaplotypeCaller for better accuracy of SNV and indel detection.

### Sample-level quality assurance (QA) checks

Three quality assurance checks were applied prior to joint genotype calling to identify problematic samples:

1. *SNV concordance check* with existing SNP array genotypes to identify possible sample errors. Using verifyBamID (Jun et al. 2012) to compare between SNP array data and WES BAM files, samples with concordance <0.95 were excluded.
2. *Sex check* for variants outside PAR region to identify possible sample swaps or misreporting. Using PLINK, samples with F statistics < 0.2 or > 0.8.
3. *Contamination check* for possible sample swaps. Using verifyBamID (Jun et al. 2012) to calculate the concordance estimate between the array genotypes and the GRCh38-mapped BAM file. A sample is potentially contaminated if the CHIPMIX value is <0.05.

In total, we dropped 55 samples that failed the SNP concordance check, 41 samples that were recorded with the incorrect sex, and 211 samples that failed the contamination check. An additional 266 samples were dropped due to consent issues, resulting in a call set containing 20,504 samples.

### Joint-genotype call of 20,504 WES samples

All WES samples were joint-called using GATK4.1.1 to create a joint-genotype called, project-level VCF (pVCF). This included four major steps:

- CombineGVCF and GenotypeGVCF – these two steps are the same in both VCPA-WGS and VCPA-WES pipelines. gVCFs of all 20,504 samples were combined in parallel across 5,000 genomic windows/regions across all the chromosomes.
- Generating the VQSR model – a Variant Quality Score Recalibration (VQSR) indicator is used for defining qualities of variants via a machine-learning model. Only variants that were called within any of the capture kits were used to build the VQSR model.
- Applying the VQSR model – the trained VQSR model was applied to all the autosomal chromosomes, as well as chromosomes X and Y, and mitochondria.

### Quality control (QC) protocol for WES samples

The GCAD quality control (QC) pipeline uses a modified protocol originally developed by the ADSP QC Working Group on WGS (Naj et al. 2019) and includes several major components: (1) pre-QC quality checks; (2) variant-level QC; (3) sample-level QC; and (4) post-QC quality checks. These steps are applied to both SNVs and indels.

We implemented variant-level QC to SNVs and indels in the project-level VCFs. Data were stratified into sequencing subsets based on the capture kit, sequencing assay, and sequencing center. We applied filters in the following order within sequencing subsets, resulting in the exclusion of: (1) variants outside of designated capture regions specific to the capture kit used on a sample; (2) variants failing GATK quality assessment (those without “PASS” or in a VQSR Tranche of 99.5% or more extreme); monomorphic variants; (3) variants with a high missingness rate (≥20%); (4) variants with excessive heterozygosity and (5) variants with high average read depth (>500x). Additionally, we estimated within subsets the allelic read ratio for each variant, and within each population (race/ethnicity). We also estimated the departure from Hardy-Weinberg Equilibrium (HWE) or excess heterozygosity, which may be used as potential exclusion criteria by end-users of the data.

We also explored sample-level QC criteria and evaluated multiple filters to further exclude potential low-quality samples. We estimated multiple quality metrics within each sample including (1) counts of singleton/doubleton variant calls (to identify an excess of private variants); (2) genotype missingness rate within the sample; (3) Transition/Transversion (Ti/Tv) ratio (for SNVs only); (4) heterozygosity-to-homozygosity ratio across all variants within individuals; and (5) the mean within-sample read depth. Samples were considered for exclusion if their values for any of these criteria were greater than 6 SD from the mean value.

### Genotype concordance analyses with the previously published ADSP-Discovery WES data

Genotype calls generated by VCPA in this data set (20,504 samples) include samples that were part of the ADSP-Discovery data set (Bis et al. 2020). To provide a comparison of genotype quality, we examined the concordance between genotype calls using the ATLAS approach (Challis et al. 2012) on 10,786 samples that overlapped between the two sets. The previous genotype calling was conducted based on GRCh37, which were lifted to GRCh38 using liftOver (Kuhn, Haussler, and James Kent 2013). For these analyses, genotype concordance was defined as identical genotype calls (including missing genotypes) between the two call sets. Concordance was calculated by sample, by variant, and overall. Because VCPA employs a joint calling approach, we also investigated the impact of the capture kit coverage on genotype concordance under the hypothesis that limited coverage in additional samples of the current data set could alter the quality control metrics.

### Annotation protocol for WES samples

Variants were annotated using our published annotation pipeline with updated resources (VEP 98 (McLaren et al. 2016), CADDv1.4 (Rentzsch et al. 2019), SnpEffv4.1k (Cingolani et al. 2012)) in GRCh38, described elsewhere (Butkiewicz et al. 2018). Briefly, we assign a “most damaging consequence” via a custom prioritization routine that down-weights non-coding transcripts or transcripts flagged as undergoing nonsense-mediated decay.

## Results

### Characteristics of Capture kits

As summarized in Table 1, a total of ten capture kits were used for sequencing 20,504 individuals. Since these files were in different genome builds and file formats, GCAD first standardized them in the same file format and normalized them to the same reference genome build GRCh38. Table 3 contains additional key information about the capture kit contents, including the size of targeted genomic regions, the percentage of capture regions that are within Ensembl v94 exons, and the percentage of the Ensembl v94 exons (with flanking bp) that are captured by each of the kits.

**Table 3:** Characterization of genomic regions (in GRCh38) by each capture kit.

| Capture kit info | Number of bp | % of capture regions that are Ensembl v94 exons | % of Ensembl v94 exons (with flanking 7bp) that are captured |
| --- | --- | --- | --- |
| Agilent WES v3 capture region | 52,941,356 | 92.54 | 95.13 |
| Agilent WES v4 capture region | 53,791,945 | 92.11 | 94.14 |
| Agilent WES v5 capture region | 53,309,273 | 89.97 | 96.27 |
| Agilent WES v6 capture region | 61,549,327 | 87.78 | 95.88 |
| IDT xGen Exome Whole Exome Research Panel v1.0 w/Custom Spike-In Baits | 39,968,963 | 94.05 | 95.20 |
| Illumina Rapid Capture Exome (ICE) kit | 40,653,755 | 91.17 | 98.03 |
| Nimblegen VCRome sequencing w/Custom Spike-In Baits | 41,708,836 | 96.26 | 93.63 |
| Roche Nimblegen's VCRome v2.1 | 37,951,907 | 96.05 | 94.39 |
| Roche SeqCap EZ Exome Probes v2.0 Target Enrichment Probes | 47,294,869 | 94.68 | 92.34 |
| Roche SeqCap EZ Exome Probes v3.0 Target Enrichment Probes | 69,167,416 | 83.01 | 97.14 |

We observed a wide range of differences in terms of the bases covered by each of these capture kits with respect to the human reference genome (from 37 million to 69 million bps). An average of 91.76% of the capture regions per capture kit were annotated as Ensembl exons. In addition, on average 95.22% of these exons were captured by each capture kit.

### Capture kit comparison

We next compare the target region designs among different capture kits. First, the Jaccard similarity measure was calculated on all capture regions at basepair level across these kits. A value of 1 indicates that the kits are very similar to each other, while a 0 indicates the opposite. The results are visualized in Figure 1.

**Figure 1:**
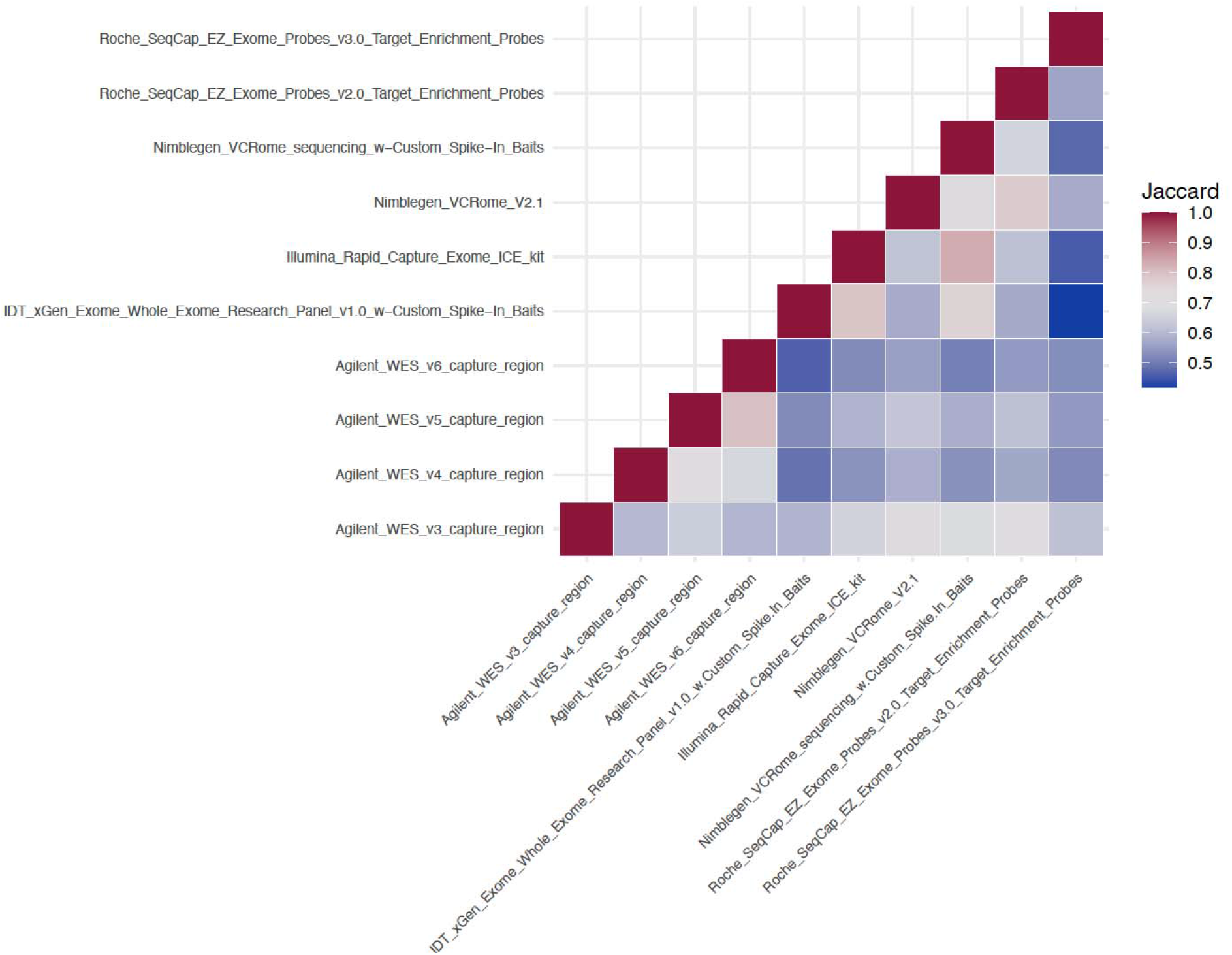
Jaccard similarity measure of the capture kits. The average of all pairwise Jaccard similarity scores is 0.586 (SD: 0.038). The two most similar kits are “Illumina_Rapid_Capture_Exome_ICE_kit” and “Nimblegen_VCRome_sequencing_w-Custom_Spike-In_Baits” (Jaccard score = 0.83). Conversely, the two most dissimilar kits are “IDT_xGen_Exome_Whole_Exome_Research_Panel_v1.0_w-Custom_Spike-In_Baits” and “Roche_SeqCap_EZ_Exome_Probes_v3.0_Target_Enrichment_Probes (Jaccard score = 0.39).”

### Data quality – WES CRAMs

The VCPA pipeline generated all CRAMs without using any capture kit information. Therefore, the differences observed in the CRAM metrics are independent of the capture kits and are primarily due to differences in the sequencing platforms used by the different sequencing centers.

We investigated whether the processed CRAMs were affected by sequencing centers (Figure 2a) or platforms (Figure 2b). We compared multiple CRAM metrics, including i) percentage of mapped reads; ii) percentage of duplicated reads; iii) percentage of paired reads; and iv) quality of reads (based on a Q score of 30 [Q30]).

**Figure 2:**
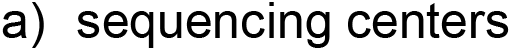
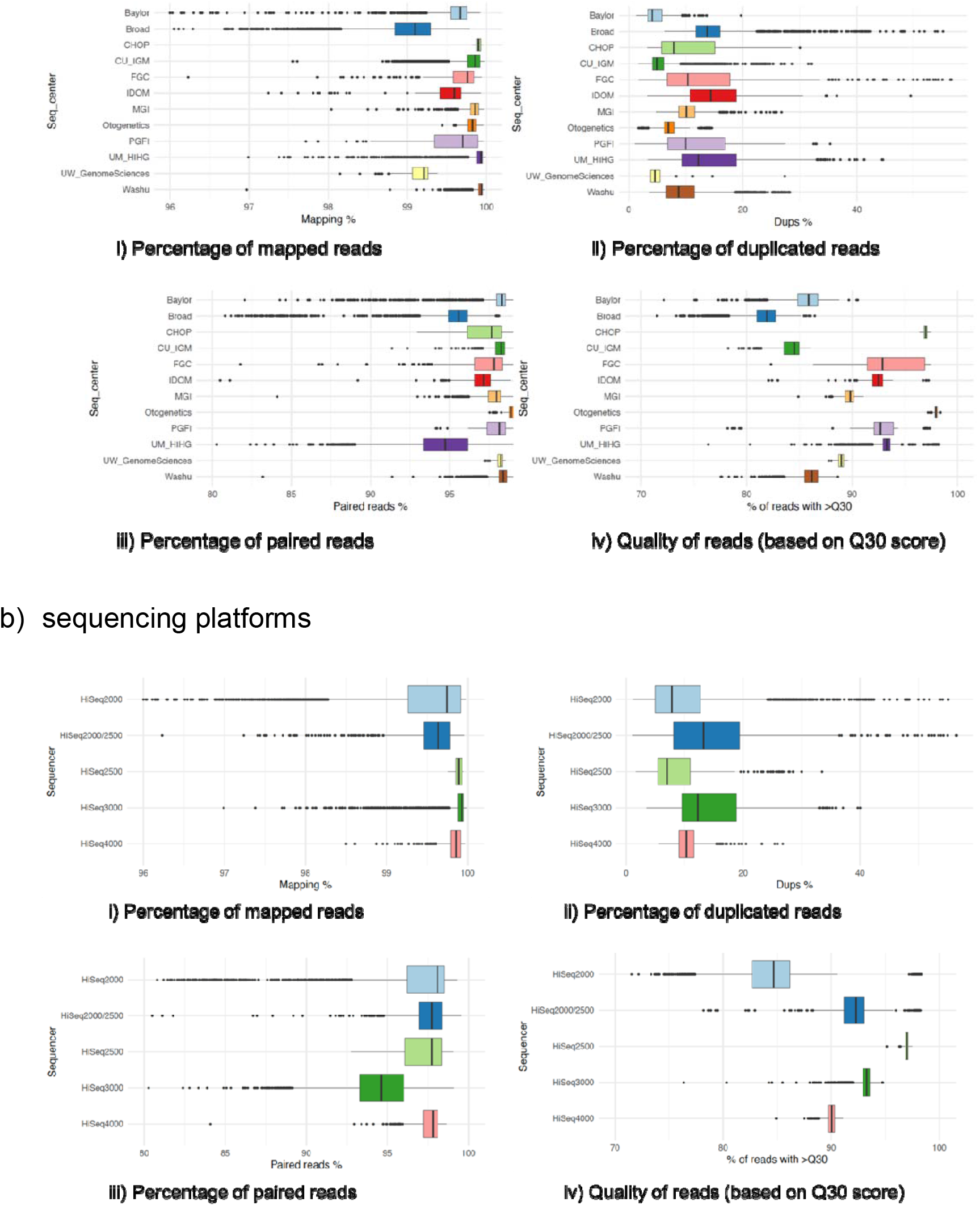
Comparison of WES CRAM quality metrics across a) sequencing centers (Seq_center); and b) sequencing platforms (Sequencer).

From Figure 2a, we observed that (1) the average mapping rate is 99.6%; (2) 93.4% of samples have <20% duplicated reads; (3) 83.4% of samples have >95% proper pairs; and (4) 97.2% of samples have >80% of alignment with Q30. While there is some variability in these metrics, we do not observe systematic bias as to which sequencing center performed best/worst in all areas as compared to the others (Figures 2a, 2b).

Next, we sought to compare the 20x coverage (i.e., percentage of bps with 20 reads or more within the sample-specific capture regions) across all samples (Figure 3). 20x coverage was chosen as it was the minimum coverage required to successfully genotype 95% of heterozygous SNPs in an analyses (Meynert et al. 2014). For about 95% of the CRAMs, we observed that >80% of reads were located within the capture region at 20x coverage. Similarly, this metric does not have sequencing center- or sequencer-specific effects, but on average, 20x coverage is lower for samples sequenced using the Illumina 2000/2500 platform.

**Figure 3:**
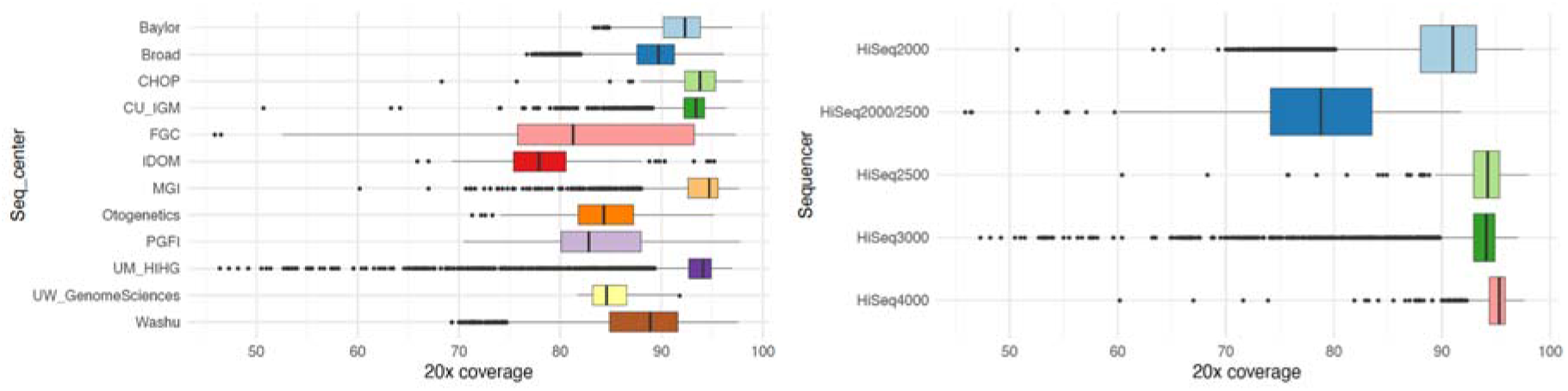
Comparison of 20x coverage of all the WES BAMs/CRAMs by a) Sequencing centers (left); and b) Sequencers (right).

### Data quality – variants

Next, we examined the variant-level data quality. GATK outputs VQSR scores. Overall, 96.83% of the variants (>7.3 million SNVs and 0.61 million indels) were labeled PASS by the model.

Besides using the GATK VQSR indicator to specify the quality of a variant, the ADSP/GCAD QC pipeline (Naj et al. 2019) outputs a series of quality metrics to determine whether the variant is within capture regions, the call rate, depth, and Ti/Tv ratio after QC. Figure 4 shows the Ti/Tv ratio on the exonic variants (colored by study) before and after QC (Before QC on the x-axis, After QC on the y-axis). Before QC, the average Ti/Tv ratio is 2.53. After QC, the Ti/Tv ratio on exonic regions in our studies is around 3.03. This post-QC Ti/Tv ratio is similar to reports in previous findings (Lek et al. 2016).

**Figure 4:**
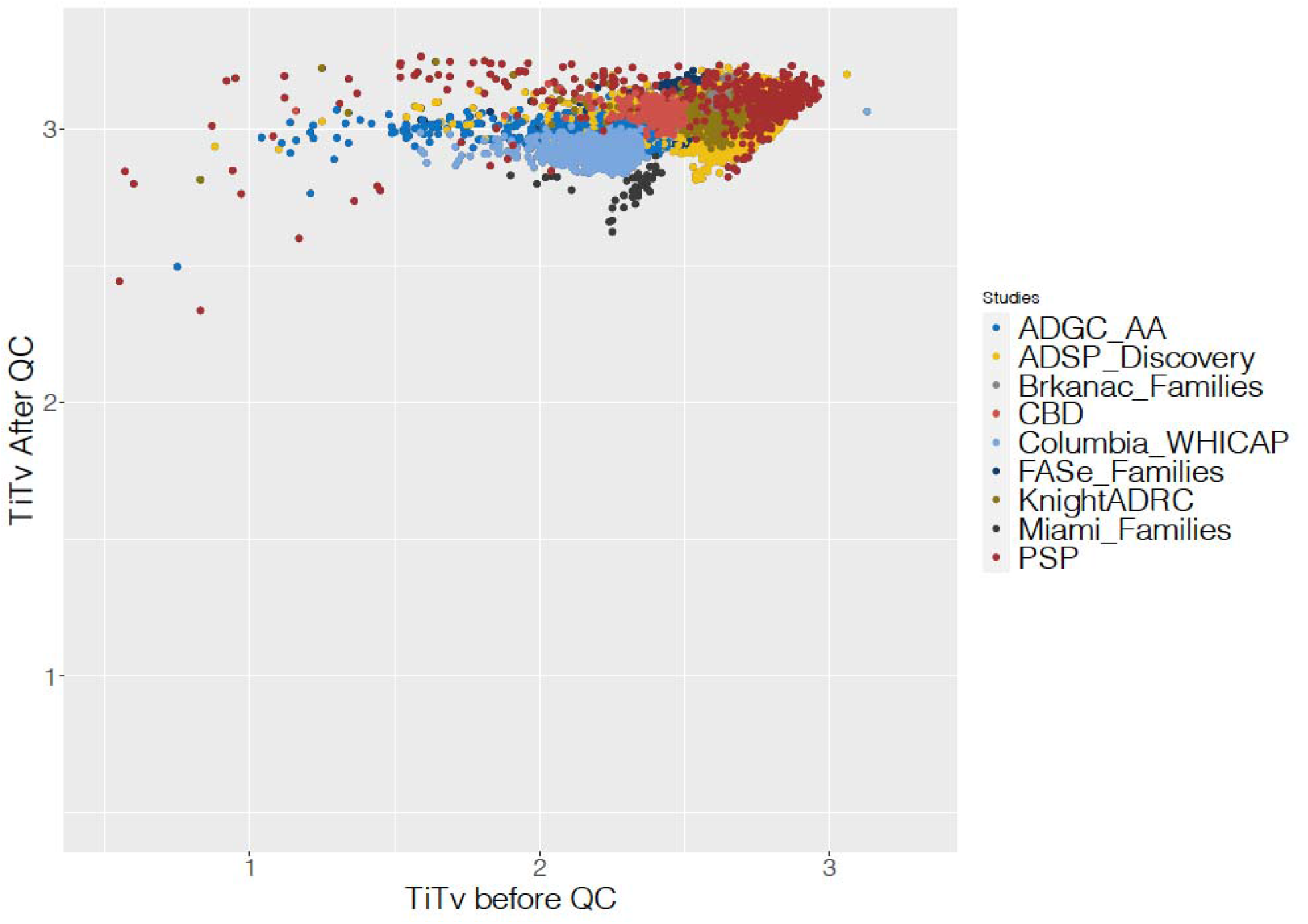
Comparison of the Ti/Tv ratio of exonic variants before and after QC. The QC protocol enables us to look for variants found across studies as well as those which are study-specific. On average, 97.26% of variants have a GATK PASS across study-capture combinations. 91.45% of variants within the designed capture kit per each study-capture combination are labeled as good quality.

### Genotype Concordance between two different callers on a set of overlapping individuals

The ADSP-Discovery data set, comprised of 10,786 individuals, was sequenced and processed by three sequencing centers: Broad Institute, Baylor College of Medicine’s Human Genome Sequencing Center, and Washington University’s McDonnell Genome Institute. Genotypes for bi-allelic SNVs and indels were called using ATLAS2 on hg19/GRCh37 (Bis et al. 2020). To evaluate the genotype quality on our 20k WES call set, which was generated using a novel approach in which no capture regions were used for individual sample calling, we examined the overall concordance, by sample and by variant, of genotypes called differently on the 10,786 samples that were present in both the ADSP-Discovery data set and the current data set. 1,407,006 variants were called in both sets, comprising 15,175,966,716 genotypes. Overall concordance was 99.43%. There were five samples with a genotype concordance <95%. Three samples had extremely low concordance (8.21%, 10.44%, and 23.44%), reflecting low DNA concentration samples, and the other two samples had 86.4% and 90.5% concordance. We also examined variant-level genotype concordance relative to capture kit coverage. The majority (69.3%) of variants were covered by all ten capture kits, 18.1% by nine, 7% by eight, and 4% by seven capture kits. These patterns may be due to differences in sequence read coverage from the various capture kits combined with joint calling approaches that leverage information across samples, but this trend only holds for a very small percentage of all called variants.

### Annotation results

Using the ADSP annotation pipeline (Butkiewicz et al. 2018), we found that the 8.16 million variants are located in 28,579 transcripts based on Ensembl annotations (McLaren et al. 2016; Butkiewicz et al. 2018). Every variant is annotated based on the most damaging VEP predicted consequence (see “Annotation protocol for WES samples” section). Figure 5A shows the top ten most damaging consequence categories. In summary, 41% of the variants are intronic, 15% are missense variants, followed by upstream/downstream gene variants (each 9%), synonymous variants (8%), and 3’UTR variants (7%). Figure 5B shows the proportion of the CADD score that is PHRED-like scores ranging from 1 to 99, based on the rank of each variant relative to all possible 8.6 billion substitutions in the human reference genome. The mean CADD score of all variants is 9.26, while the median value is 6. Meanwhile, 15.5% of the variants have a CADD score >20, meaning that these variants are among the top 1% of deleterious variants in the human genome. In addition, 1.8% of the WES variants are among the top 0.1% of deleterious variants in the human genome (CADD>30).

**Figure 5:**
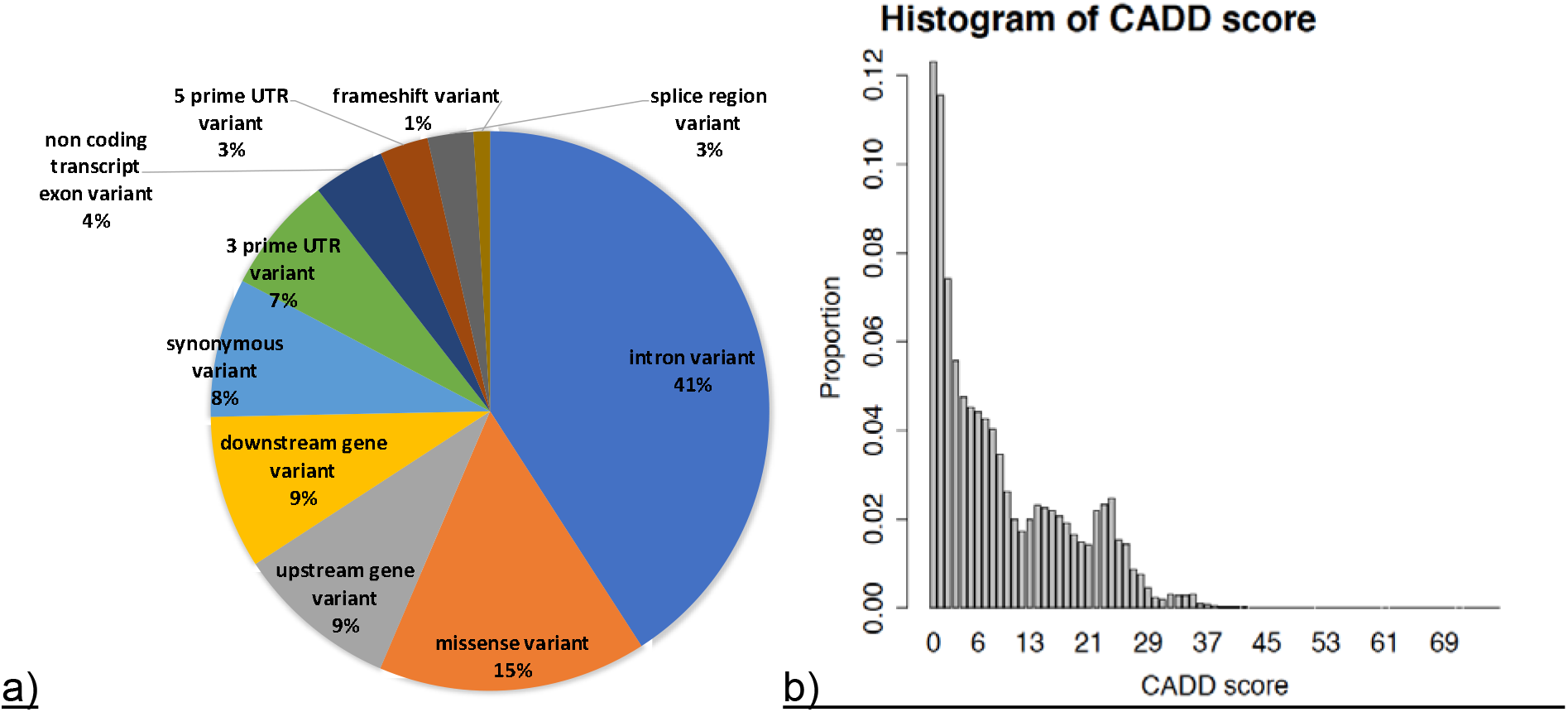
Annotation results for all variants called in the WES pVCF of 20,504 samples: a) Top ten categories of VEP predicted consequence; b) Distribution of CADD phred-normalized scores.

### Data sharing – NIAGADS Data Sharing Service (DSS)

The National Institute on Aging Genetics of Alzheimer’s Disease Data Storage Site (NIAGADS) is a national data repository that facilitates access to genetic data by qualified investigators for the study of the genetics of early-onset/late-onset Alzheimer’s Disease and Alzheimer’s Disease Related Dementias (ADRD). Collaborations with large consortia such as the Alzheimer’s Disease Genetics Consortium (ADGC), Cohorts for Heart and Aging Research in Genomic Epidemiology (CHARGE) Consortium and the ADSP, a main mission of NIAGADS is to manage large AD genetic data sets that can be easily accessed by the research community.

The NIAGADS Data Sharing Service (DSS) released the CRAMs (compressed version of BAM files), gVCFs generated by GATK4.1.1, and QC-ed pVCFs of the abovementioned ADSP WES data set in September 2020 (NG00067.v3), together with pedigree structures for family studies and phenotypes that were harmonized according to ADSP protocols. Qualified investigators can access these data with a submission request and approval from the NIAGADS Data Access Committee managed by independent NIH program officers. Data can be downloaded through the DSS portal. More information about the data set can be found on the data set page,◻NG00067 (https://dss.niagads.org/datasets/ng00067/). See the◻Application Instructions◻page (https://dss.niagads.org/documentation/applying-for-data/application-instructions/) on how to submit a Data Access Request and access data.

## Discussion and conclusions

In this study, we developed a new bioinformatics approach to joint call WES samples sequenced using multiple capture kits from different sequencing providers. The procedure has been successfully applied to a total of 20,504 exomes gathered through the collaborative network of the ADSP, resulting in the generation of the world’s largest publicly available AD WES data collection and joint-call pVCF to date. Annotated and QC-ed following ADSP protocols, this high-quality WES pVCF with ~7.5 million SNVs and >700,000 indels is publicly available at NIAGADS DSS for qualified investigators worldwide.

Joint-calling WES samples based on different capture kits poses several challenges due to the nature of the kits. Although each capture kit was designed primarily based on exonic regions (> 91% of the capture regions per capture kit were covered by Ensembl exons, (**Table 3**, **Section Characteristics of Capture kits**), they were also designed based on different genome builds and gene annotations, therefore resulting in substantial differences in the captured contents (average Jaccard similarity score of ~0.6 per capture kit vs all other kits, (**Figure 1**, **Section Capture kits comparison**). In order to successfully harmonize/joint-call all of the data without any systematic bias, a uniform bioinformatics pipeline together with the standardization of capture kit target region definitions are critical.

In our calling strategy, we first lifted over all capture kits using the same protocol to GRCh38 if they were not of this genome build. We then processed all WES samples using a single analysis pipeline: VCPA-WES (**Section VCPA for WES processing**). We did not use capture region definitions to limit variant calls when we generated gVCF or joint-called pVCFs. Capture region definitions were only used in the QC steps to identify high quality variant and genotype calls. There are two advantages of this approach. First, in the future when additional samples on different capture kits needs to be incorporated, we can reuse the gVCFs of these old samples without reprocessing gVCFs. Second, we can retain variants and genotype calls that are either outside the target regions but still have good quality, or in regions that are not targeted by all capture kits. Indeed, if we only look at regions that are targeted by all capture kits, we are left with only about 45% of the unions.

The genotype data generated by this tailor-made bioinformatics strategy are high-quality. We have shown in Figures 2 and 3 that there is no systematic bias attributed to the sequencing centers and sequencers on the CRAM quality (**Section Data quality – WES CRAMs**). This shows that even though the WES data were sequenced in different ways, these experimental artifacts could be greatly reduced with a carefully designed data processing pipeline.

Next, we evaluated the quality of the variants using the GATK VQSR score, Ti/Tv ratio (Figure 4), and via our in-house GCAD/ADSP QC protocol (**Section Data quality – variants**). Overall, ~97% of variants were labeled PASS by GATK. After variant-level QC was performed, the Ti/Tv ratio in exonic regions in our studies was >3. This is similar to what was previous reported, indicating that the data doesn’t contain many false positives caused by random sequencing errors. In terms of study-capture specific variants, using a multi-step QC protocol as shown in Table 4 (exclude variants with high missingness rate, excessive heterozygosity, high read depth, etc.), >91% of variants have successfully met our QC criteria, are of good quality, and can be used in subsequent analyses.

**Table 4:** Variant quality per study-capture combination.

| Study | Capture Kit info | % of variants that are monomorphic | % of non-monomorphic sites, in capture kits | Within the capture kits, with GATK PASS % | Within the capture kits, with VFLAG 0 % |
| --- | --- | --- | --- | --- | --- |
| MIAMI family | Agilent WES v3 capture region | 97.65% | 63.22% | 97.56% | 91.70% |
| MIAMI family | Agilent WES v4 capture region | 97.88% | 57.95% | 95.29% | 72.50% |
| CBD | Agilent WES v5 capture region | 94.20% | 68.74% | 98.09% | 95.30% |
| FASe family | Agilent WES v5 capture region | 93.60% | 70.88% | 97.57% | 91.84% |
| ADGCAA | Agilent WES v6 capture region | 71.92% | 73.71% | 97.56% | 94.81% |
| Knight ADRC | IDX xGen Exome Whole Exome Research Panel v1.0 w/Custom Spike-in Baits | 95.90% | 47.79% | 97.44% | 95.48% |
| ADSP-Discovery | Illumina Rapid Capture Exome (ICE) kit | 77.87% | 54.20% | 97.90% | 93.87% |
| FASe family | Nimblegen VCRome sequencing w/Custom Spike-in Baits | 96.06% | 46.57% | 96.73% | 93.86% |
| Knight ADRC | Nimblegen VCRome sequencing w/Custom Spike-in Baits | 91.67% | 47.97% | 97.56% | 95.31% |
| ADSP-Discovery | Roche Nimblegen's VCRome v2.1 | 69.64% | 66.22% | 98.51% | 94.69% |
| FASe family | Roche Nimblegen's VCRome v2.1 | 95.91% | 62.59% | 96.95% | 91.74% |
| PSP | Roche Nimblegen's VCRome v2.1 | 92.47% | 59.10% | 95.60% | 84.13% |
| Brkanac | Roche SeqCap EZ Exome Probes v3.0 Target Enrichment Probes | 97.17% | 64.59% | 97.45% | 91.17% |
| Columbia WHICAP | Roche SeqCap EZ Exome Probes v3.0 Target Enrichment Probes | 67.19% | 78.37% | 97.39% | 93.90% |

Lastly, we performed genotype concordance analyses on the set of overlapping samples found in both the previously published ADSP-Discovery WES data set and this newly joint-called WES data set. Around 11,000 samples were used for the analyses (**Section Genotype Concordance between two different callers on a set of overlapping samples**). Even though the two data sets were called using different callers (ATLAS vs GATK), >1.4 million variants were called in both data sets, comprising 15 billion genotypes. Overall concordance was 99.43%. Variants that were not concordant may be located in genomic regions that are difficult to be sequenced. This shows that, despite the complexity involved in creating this new, much larger, joint-called WES data set, the innovative bioinformatics strategy allowed us to produce a data set with high-quality genotypes.

The 8.16 million variants in this WES data set span across 28,579 transcripts (**Section Annotation results**). We annotated every variant based on the most damaging VEP predicted consequence (Figure 5A). The top ten most damaging consequence categories included missense variants, upstream/downstream gene variants, synonymous variants, and 3’UTR variants. Meanwhile, 15.5% of the variants have a normalized CADD score >20, meaning that these variants are among the top 1% of deleterious variants in the human genome (Figure 5B). These results showcase that the ADSP annotation pipeline we developed is very well capable of annotating both WGS and WES data.

In conclusion, the VCPA-WES bioinformatics pipeline, together with the QC and annotation protocol GCAD developed, enable us to generate a high-quality AD-specific WES data set containing over 8 million variants on 20,504 samples. The pipeline works well when joint-calling any WES data (of different phenotypes) that are sequenced in different batches using different sequencing machines and capture kits. This valuable data set is free of batch effects and is, by far, the largest publicly available AD WES data set. It is available at NIAGADS DSS: https://dss.niagads.org/datasets/ng00067/. Qualified and approved investigators can apply to access and download the data for various research purposes.

## Author contributions (scientific reports requirement)

Y.Y.L., A.C.N., Y.-F.C., and W.S.B. conceived and designed the experiments. Y.Y.L., Y.-F.C., N.W. and W.S.B. performed data analyses. Y.-F. C., O.V., P.G., L.Q. carried out the data production under the supervision of Y.Y.L., A.C.N. and E.M. A.B.K., L.C. and H.I. manages phenotypes for these samples. Y.Y.L., H.L., K.C., W.-P. L., and L.-S.W. wrote and edited the manuscript. G.D.S., L.-S. W., E. M., A.C.N. and A.D. secured funding. All authors read and approved the manuscript.

## Data availability

All CRAMs (compressed version of BAM files), gVCFs generated by GATK4.1.1, and QC-ed pVCFs of the abovementioned ADSP WES data set are available in the NIAGADS Data Sharing Service (DSS) (NG00067.v3), together with pedigree structures for family studies and phenotypes that were harmonized according to ADSP protocols. Qualified investigators can access these data with a submission request and approval from the NIAGADS Data Access Committee managed by independent NIH program officers. Data can be downloaded through the DSS portal. More information about the data set can be found on the data set page,◻NG00067 (https://dss.niagads.org/datasets/ng00067/). See the◻Application Instructions◻page (https://dss.niagads.org/documentation/applying-for-data/application-instructions/) on how to submit a Data Access Request and access data.

